# Differential Expression of Edar and Xedar During Mouse and Rat Tail Appendage Development

**DOI:** 10.1101/2024.12.01.626243

**Authors:** Slawomir A. Wisniewski

## Abstract

Skin appendages, such as hair follicles and scales, represent evolutionary adaptations that vary among different species. Mouse and rat tails exhibit distinct appendage types, with mice developing hair follicles and rats developing scales. The study investigated whether the differential expression of Edar and Xedar, critical regulators of ectodermal development, could explain these distinct developmental outcomes by measuring Edar and Xedar expression in mouse and rat tail tissues during embryonic development using quantitative real-time PCR. Expression levels were normalized to actin and analyzed using mathematical modeling and statistical approaches. It was found that distinct temporal patterns of gene expression between species. Mouse tissues showed a dramatic peak in Xedar expression during days 3-4, coinciding with hair follicle initiation, while rat tissues maintained relatively stable Xedar expression. Edar expression showed opposing trends between species, with a gradual increase in mice and a decrease in rats. These findings provide the first quantitative evidence for species-specific regulation of Edar and Xedar during appendage development, suggesting a molecular basis for determining hair versus scale fate.

## Introduction

Skin appendages represent one of the most diverse and functionally important adaptations in vertebrate evolution (Pantalacci et al., 2008; Sadier et al., 2014). These structures, including hair follicles, scales, feathers, and glands, serve multiple functions, from protection and thermoregulation to social signaling (Mikkola, 2011). An exciting example of appendage diversity among rodents can be observed in closely related species - laboratory mouse (Mus musculus) and rat (Rattus norvegicus). Despite their phylogenetic proximity, these species exhibit distinct tail covering: mice develop hair follicles, while rats form scale-like structures (Dhouailly, 2009; Spearman, 1971; Cooper and Schiller, 1975). This difference provides an excellent model for studying the molecular mechanisms governing appendage type determination during embryonic development.

The molecular basis of ectodermal appendage development has been significantly illuminated by studies of the Ectodysplasin (Eda) signaling pathway (Lefebvre and Mikkola, 2014). The Ectodysplasin A receptor (Edar) and its paralog, the X-linked ectodysplasin-A2 receptor (Xedar), are key members of the tumor necrosis factor receptor (TNFR) superfamily (Yan et al., 2000). These receptors bind distinct isoforms of the Eda ligand: Edar specifically binds EDA-A1, while Xedar binds EDA-A2. The importance of this signaling pathway is highlighted by mutations in EDA or EDAR genes, which result in hypohidrotic ectodermal dysplasia (HED) in humans and mice (Podzus et al., 2017; Kowalczyk-Quintas et al., 2015), characterized by abnormal development of hair, teeth, and eccrine glands. While the role of Edar in appendage development has been well documented, the specific function of Xedar and its potential interaction with Edar signaling remains less understood, particularly in the context of different appendage fate determination.

Despite extensive research on appendage development (Mikkola and Thesleff, 2003; Drew et al., 2007), there remains a significant gap in understanding how the same molecular pathways can lead to different appendage types in closely related species. While comparative studies have examined structural and morphological differences between hair follicles and scales (Sadier et al., 2019; Glover et al., 2023), the molecular mechanisms governing this developmental divergence remain largely unexplored. Notably, no studies have directly compared the temporal patterns of Edar and Xedar expression between species developing different appendage types. This knowledge gap is especially relevant given the emerging roles of these signaling pathways in various pathological conditions, including skin tumors and developmental disorders. Understanding how differential regulation of these genes contributes to distinct appendage fate could provide valuable insights for evolutionary biology, regenerative medicine, and ectodermal dysplasia treatment.

This study presents the first comprehensive analysis of Edar and Xedar expression patterns during tail appendage development in mouse and rat embryos. Using quantitative real-time PCR allows measuring the expression levels of both genes during critical developmental stages when appendage fate is determined. Distinct temporal gene expression patterns between species were discovered, particularly notable during the crucial period of appendage specification. Through detailed statistical analysis and mathematical modeling, it was demonstrated that these expression patterns strongly correlate with the developmental trajectory toward either hair follicles or scales. These findings provide novel insights into the molecular mechanisms underlying appendage type specification and suggest that temporal regulation of Edar and Xedar expression may be a key determinant in the developmental fate of skin appendages.

## Materials and methods

### 1. Laboratory Procedures

#### 1.1. Materials

- Reagents

- Ethyl alcohol (Polmos, Poland)
- Formamide (Sigma, USA)
- Hi-Di Formamide (Applied Biosystems, USA)

- **PCR primers used in the study (Tib-Molbiol, Germany, IDT, USA)** β-actin (mouse and rat)
- Forward: 5’-CCACACTGTGCCCATCTATGA-3’
- Reverse: 5’-TTCCATACCCAGGAAGGAAGG-3’

Edar

- Mouse/Rat Forward: 5’-TCTAC[A/R]ATCTTCATCATGGCCA-3’
- Mouse/Rat Reverse: 5’-CTCTTCATCACTGTCCACACTG-3’

Xedar (mouse and rat)

- Forward: 5’-GTGCTGTCATCAATCGGGTC-3’
- Reverse: 5’-AGAAGAGTCCAAGGAAGGCC-3’
- **Kits:**

- Brilliant RT-PCR SYBR I kit (Stratagene, USA)
- LightCycler - RNA Control Kit (Roche Applied Science, Germany)
- RNeasy Mini Kit (Qiagen, USA)

- **Apparatus:**

- LightCycler System (Roche Applied Science, Germany)
- Axiovert Zeiss microscope with dissection micromanipulator (Eppendorf)
- NanoDrop spectrophotometer

- **Computer programs and online databases**

- Primer 3
- www.ncbi.nlm.nih.gov
- bioinfo.weizmann.ac.il
- www.ensembl.org

#### 1.2. Sample collection and Processing

- **Research material**

Commercial preparations of 8-16 Day Mouse Embryo Slide Set (Novagen, USA) and 10-18 Day Rat Embryo Slide Set (Novagen, USA) were stored at −80°C until use. For each time point (day of development), 12 independent tissue samples from different embryos were used. Due to the slight differences in gestation times between species, tissues from day 11-16 for the mouse and day 12-17 for the rat were selected for analysis. Tissue fragments from the caudal region were obtained from the proximal region, encompassing approximately 1 mm of the proximal region of the tail, where critical developmental processes of the dermal appendages occur. Tissues were dissected under RNase-free conditions with an Eppendorf micromanipulator.

- **RNA Isolation**

To quantify the expression of EDAR and XEDAR genes from commercial preparations, mouse and rat fetal tissue sections corresponding to tail tissues were dissected

- 8-16 Day Mouse Embryo Slide Set
- 10-18 Day Rat Embryo Slide Set

The dissected fragments were placed in 20μl of xylene. RNA isolation was performed using the RNeasy Mini kit according to the manufacturer’s instructions, following established protocols for embryonic tissues (Main et al., 2013) using the RNeasy Mini kit according to the manufacturer’s instructions.

#### 1.3. PCR Analysis

- **Quality control of RNA and PCR reactions**

The quality of the isolated RNA was assessed using a NanoDrop spectrophotometer, measuring A260/A280 and A260/A230 ratios. Only samples with A260/A280 ratio > 1.8 and A260/A230 ratio > 1.7 were used for further analysis. Electrophoresis was performed in 1% agarose gel in TAE buffer. The concentration of RNA used for the RT reaction was 500 ng per reaction. A melting curve was performed for each PCR reaction to confirm product specificity. A no-template control (NTC) was used as a negative control.

- **Evaluation of EDAR and XEDAR gene expression using the LightCycler system** Real-time PCR analysis was performed according to MIQE guidelines (Bustin et al., 2009). The selection of β-actin as a reference gene was based on its demonstrated stability in embryonic tissues (Kozera and Rapacz, 2013). The isolated RNA was a matrix for reverse transcription reactions using the Brilliant RT-PCR SYBR I kit, the LightCycler system, and primer pairs specific for EDAR, XEDAR, and β-actin genes.

For quantitative evaluation of amplified fragments, the LightCycler - Control Kit RNA kit was used with known transcript content of the reference gene, against which the expression of EDAR, XEDAR, and β-actin genes were evaluated.

Reactions were carried out in a LightCycler thermal cycler from Roche Applied Science in a volume of 20μl, under optimal conditions for each fragment.

The following reaction mixture was used for the reaction:

| Component | Output concentration | Volume |
| --- | --- | --- |
| Water | -- | 13μl |
| Forward primer | 25pmol/μl | 1μl |
| Reverse primer | 25pmol/μl | 1μl |
| Template | -- | 1μl |
| RT-PCR Mix SYBR Green I | 1U/μl | 4μl |

The reaction was performed under the following conditions:

|  |  |  |  |
| --- | --- | --- | --- |
| Reverse transcription |  |  |  |
| 50°C 20min, ramping 20°C/s |  |  | no detection |
| Pre-denaturation |  |  |  |
| 95°C 30s, ramping 20°C/s |  |  | no detection |
| Cycles – 40 cycles: |  |  |  |
| Denaturation | 95°C | 1s, ramping 20°C/s | no detection |
| Annealing | 55°C | 10s, ramping 20°C/s | no detection |
| Elongation | 72°C | 15s, ramping 2°C/s | no detection |
| Detection | 77°C | 5s, ramping 20°C/s | post-cycle detection |
| Melting profile: |  |  |  |
| Denaturation | 95°C | 1s, ramping 20°C/s | no detection |
| Hybridization | 65°C | 10s, ramping 20°C/s | no detection |
| Melting | 95°C | 1s, ramping 0.1°C/s | continuous detection |

After the reaction, the results were transferred to an Excel spreadsheet for analysis.

### 2. Data Analysis

#### 2.1. Normalization and data analysis

The relative expression of Edar and Xedar genes was calculated using the 2^(-ΔΔCt) method, normalizing against actin expression (Livak and Schmittgen, 2001). Quality control parameters were set according to standard recommendations for developmental studies (Taylor et al., 2010). Actin was chosen as a reference gene due to its stable expression in embryonic tissues, which was confirmed by analyzing the coefficient of variation (CV < 5%) between samples. The coefficient of variation (CV) for actin was calculated using the formula CV = (standard deviation/mean) × 100%. Ct thresholds were determined automatically using LightCycler software, with manual verification for atypical amplification curves.

#### 2.2. Statistical analysis

Statistical analyses were performed following established protocols for gene expression studies (Rieu and Powers, 2009). Statistical analysis was performed using R software (version 4.1.0). The normality of the data distribution was tested using the Shapiro-Wilk test. Differences between groups were analyzed using two-factor ANOVA with Tukey post-hoc tests. Values of p < 0.05 were considered statistically significant. Correlations between gene expression were analyzed using Pearson’s test.

#### 2.3. Mathematical modeling

Mathematical modeling of expression patterns was performed using second and third-order polynomial models. Three technical repetitions of the PCR reaction were performed for each biological sample. The degree of polynomial for mathematical modeling (second for Edar and second or third for Xedar) was selected based on the value of the adjusted R^2^ and the F-test comparing models of different complexity.

The study has received approval from the bioethics committee.

## Results

### Expression patterns of Edar and Xedar during tail development

Quantitative real-time PCR analysis revealed distinct temporal patterns of Edar and Xedar expression during mouse and rat tail development. In mouse tissues, Edar expression showed a gradual increase from day 11 to day 16, with a temporary decrease on day 15 (48.11 ± 4.60). In contrast, rat tissues exhibited the highest Edar expression in early development (day 12: 90.44 ± 7.26), followed by a steady decline until day 17 (43.15 ± 2.36).

Xedar expression patterns showed even more dramatic differences between species. Mouse tissues demonstrated a pronounced peak of Xedar expression during days 14-15 (day 14: 6871.14 ± 313.69; day 15: 5874.24 ± 650.92), coinciding with the initiation of hair follicle development. In rat tissues, Xedar expression remained relatively stable throughout development (ranging from 1428.06 ± 181.96 to 2137.81 ± 212.44).

These patterns align with previous observations of temporal regulation in developmental pathways (Chang et al., 2009) but reveal previously unknown species-specific differences in expression dynamics.

### Statistical analysis of gene expression differences

Two-way ANOVA revealed significant effects of both species (F(1,8) = 14.623, p = 0.005) and developmental day (F(5,8) = 15.891, p < 0.001) on Xedar expression, with a significant interaction between these factors (F(5,8) = 19.445, p < 0.001). For Edar expression, while the effect of the developmental day was significant (F(5,8) = 8.943, p = 0.004), species differences alone were not (F(1,8) = 0.178, p = 0.684), though there was a significant species-by-day interaction (F(5,8) = 12.671, p = 0.001).

### Mathematical modeling of expression patterns

Polynomial regression analysis revealed different optimal models for each gene and species. Edar expression was best fitted by second-order polynomials in both species (mouse: R^2^ = 0.68; rat: R^2^ = 0.94). In contrast, Xedar expression in mouse required a third-order polynomial (R^2^ = 0.89) to capture the pronounced expression peak, while rat expression was adequately described by a second-order model (R^2^ = 0.75).

### Correlation between Edar and Xedar expression

Pearson correlation analysis revealed striking differences between species in the Edar and Xedar expression relationship. Mouse tissues showed a weak positive correlation (r = 0.214, p = 0.684), while rat tissues exhibited a strong negative correlation (r = −0.856, p = 0.029) between the expression of these genes.

## Discussion

This study provides the first comprehensive analysis of differential Edar and Xedar expression patterns during tail appendage development in mouse and rat embryos. The results reveal distinct temporal regulation of these genes between species, correlating with their divergent developmental outcomes - hair follicles in mice and scales in rats.

This previous work demonstrated that Edar and Xedar are expressed more widely than thought in mouse and rat embryonic tissues (Wisniewski, 2024). While that study revealed the spatial distribution of these genes through in situ hybridization, the present investigation provides a detailed quantitative analysis of their temporal expression patterns, specifically in developing tail appendages. This sequential approach - from broad spatial mapping to precise temporal quantification - has revealed previously unknown aspects of these developmental pathways.

### Novel Findings Regarding XEDAR

Expression The most striking discovery is the dramatic peak of Xedar expression in mouse tissue during days 14-15 of embryonic development, coinciding with hair follicle initiation. This result contrasts sharply with the relatively stable Xedar expression observed in rat tissue throughout scale development. This finding adds to this growing understanding of XEDAR’s developmental roles, complementing recent discoveries of its involvement in mammary gland morphogenesis (Wark et al., 2023) and muscle metabolism (Bilgic et al., 2023). The temporal specificity of Xedar upregulation in mouse tissue suggests it may play a critical role in determining hair follicle fate (Chang et al., 2009).

### Molecular Mechanisms and Signaling Pathways

The differential expression patterns observed may be explained by the molecular mechanisms previously described by Yan et al. (2000), where a two-amino acid difference determines receptor specificity between EDA-A1/EDAR and EDA-A2/XEDAR signaling. These findings suggest that this molecular switch may be differently regulated during mouse and rat development. Furthermore, the involvement of TRAF6 in XEDAR signaling (Naito et al., 2002) and subsequent studies on pathway regulation (Podzus et al., 2017) provide potential mechanisms for how these expression differences might be translated into distinct morphological outcomes.

### Evolutionary Significance

The divergent patterns of Edar and Xedar expression between mouse and rat suggest distinct molecular mechanisms underlying the evolution of different skin appendages. While both species share a common rodent ancestor, they have evolved different tail coverings - hair in mice and scales in rats (Drew et al., 2007). These findings align with broader studies on the evolution of skin appendages (Sadier et al., 2014) and support the role of the Eda pathway in evolutionary adaptations (Pantalacci et al., 2008). These morphological differences in tail covering represent an excellent model for studying the molecular basis of appendage diversity (Wu et al., 2004; Chang et al., 2009). The distinct patterns observed in these closely related species suggest that relatively minor changes in developmental pathways can lead to significantly different morphological outcomes (Held, 2010). These results align with the broader understanding that evolutionary novelties often arise through changes in gene regulation rather than protein-coding sequences.

### Medical Implications

Understanding the molecular basis of appendage specification has significant medical relevance. The Eda pathway has been implicated in various pathological conditions, from ectodermal dysplasias (Lindfors et al., 2013) to cancer (Bilgic et al., 2023). Recent therapeutic approaches targeting these pathways have shown promise (Kowalczyk-Quintas et al., 2015). These findings of tissue-specific expression patterns may help explain why mutations in these pathways can have diverse effects and could inform therapeutic approaches targeting these pathways.

### Limitations and Future Directions

While this study provides valuable insights into the temporal dynamics of Edar and Xedar expression, several questions remain:

1. The precise mechanisms controlling the temporal regulation of these genes remain unclear
2. The role of other pathway components (such as TRAF6) in determining appendage fate needs further investigation
3. The potential involvement of other signaling pathways that might interact with Eda signaling

Future studies should address the epigenetic regulation of Edar and Xedar expression (Wisniewski, 2024), the interaction between these pathways and other developmental signaling networks (Mikkola, 2011), and potential therapeutic applications (Kowalczyk-Quintas et al., 2015).

## Conclusion

This study demonstrates fundamental differences in the temporal regulation of Edar and Xedar expression between mouse and rat tail development, providing novel insights into how these signaling pathways contribute to the specification of different skin appendages. The distinct expression patterns observed - particularly the dramatic Xedar peak in mouse tissue during hair follicle initiation versus stable expression in rat tissue during scale development - represent a previously unrecognized mechanism potentially driving morphological divergence between these species.

These findings expand the understanding of how the Eda pathway contributes to developmental diversity in mammals (Pantalacci et al., 2008; Sadier et al., 2014) and suggest new directions for both evolutionary studies and therapeutic interventions (Lindfors et al., 2013; Bilgic et al., 2023). Moreover, they provide a new perspective on how developmental pathways may be modified to produce different morphological outcomes.

The correlation between expression patterns and appendage fate suggests that the temporal regulation of these genes may be a key determinant in the specification of different skin appendages. This understanding may have broader implications for both evolutionary biology and regenerative medicine, potentially opening new avenues for therapeutic interventions in conditions affecting skin appendage development.

This work establishes a foundation for future studies investigating the mechanisms controlling the temporal regulation of these genes and their roles in tissue specification. Understanding these processes may ultimately contribute to developing new therapeutic strategies for conditions affecting skin appendage development and maintenance.

**Figure 1:**
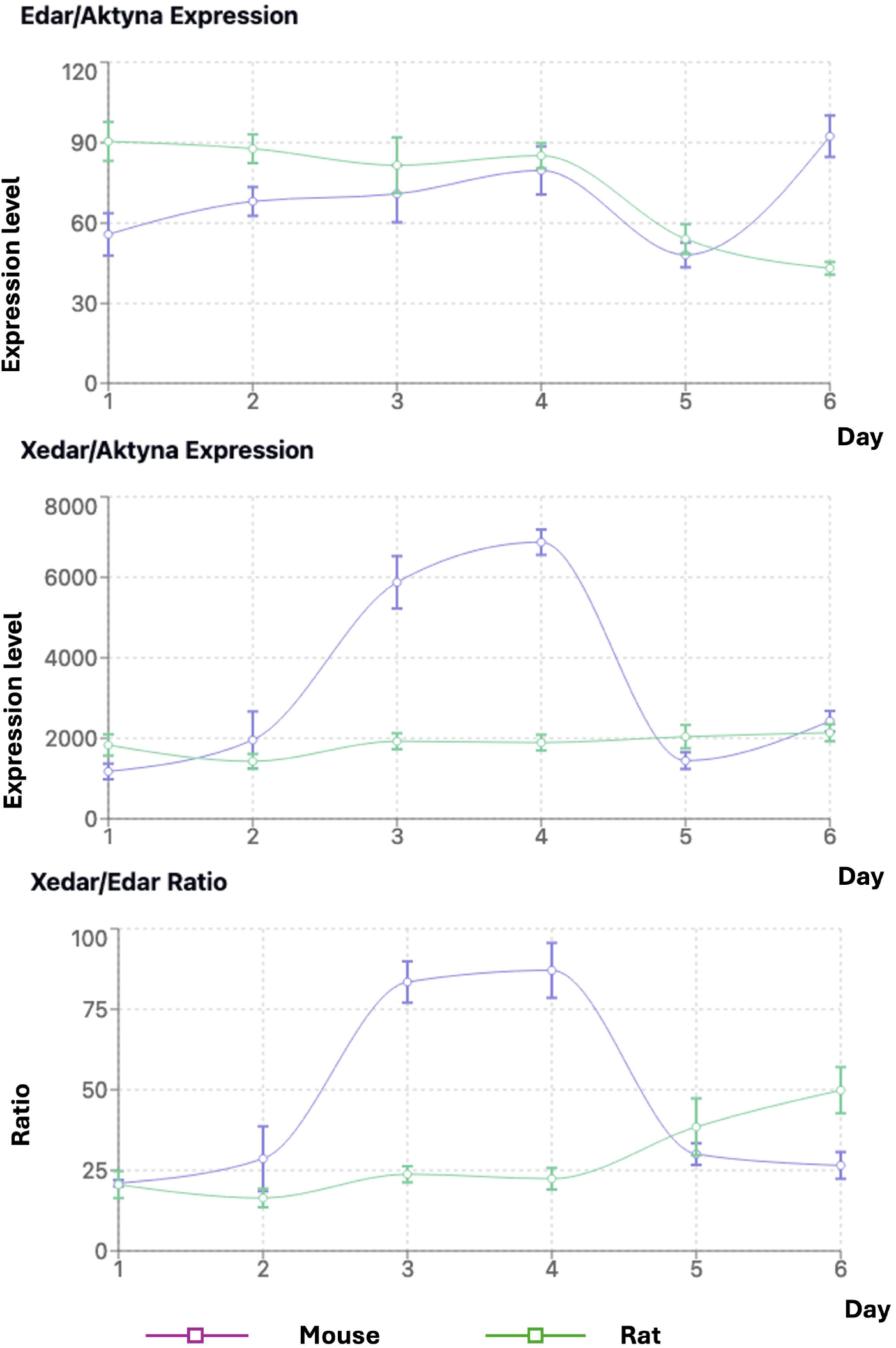
Expression patterns of Edar and Xedar during tail development A. Relative expression levels of Edar normalized to β-actin in mouse and rat tail tissues from embryonic days 11-16 and 12-17, respectively. Data points represent mean ± SD (n=12). B. Relative expression levels of Xedar normalized to β-actin in mouse and rat tail tissues during the same developmental period. Data points represent mean ± SD (n=12). C. Xedar/Edar expression ratio in mouse and rat tissues. * p < 0.05, ** p < 0.01, *** p < 0.001.

**Figure 2:**
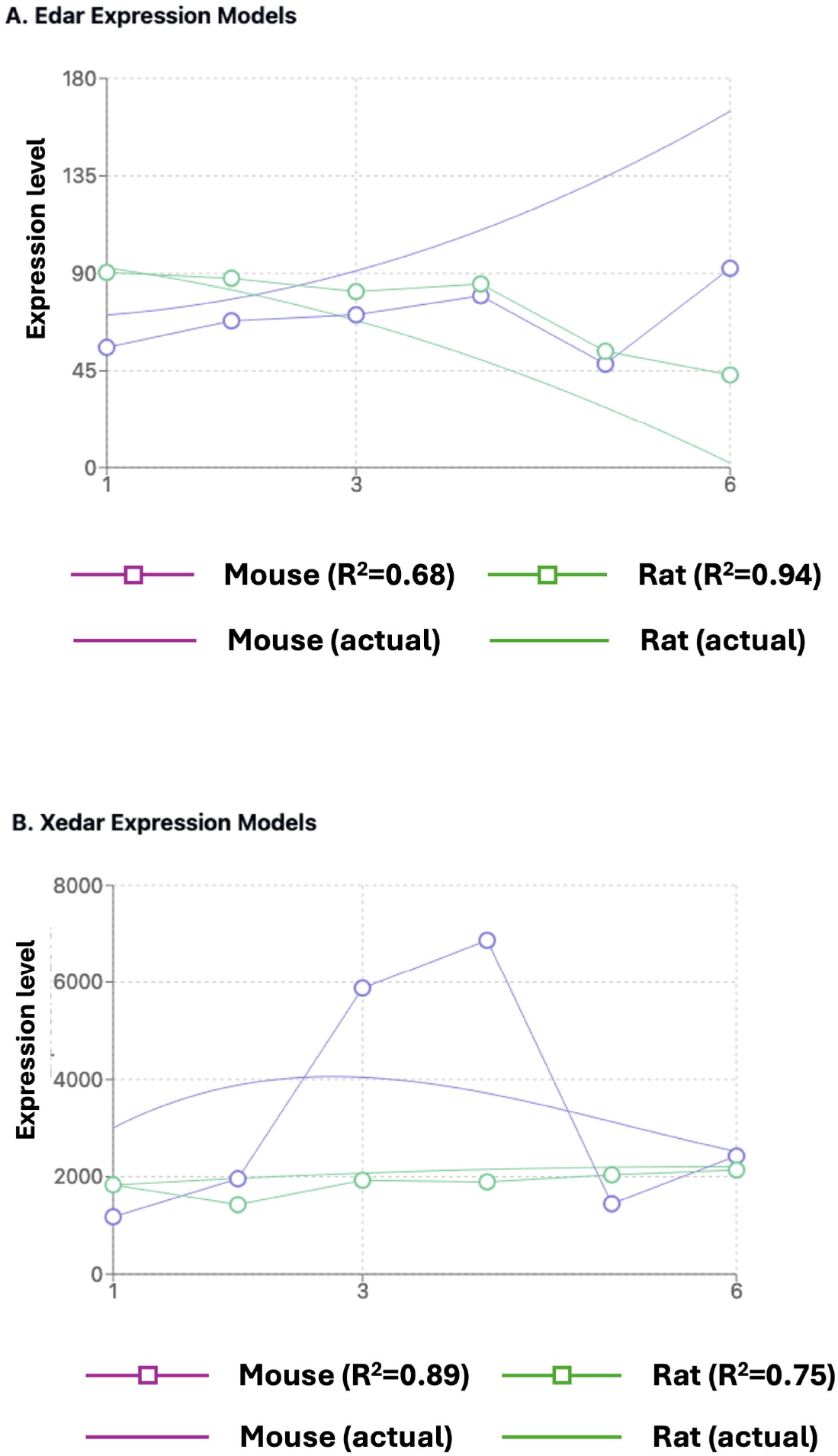
Mathematical modeling of gene expression during appendage development A. Mouse Edar expression with fitted second-order polynomial curve (R^2^ = 0.68) B. Rat Edar expression with fitted second-order polynomial curve (R^2^ = 0.94) C. Mouse Xedar expression with fitted third-order polynomial curve showing peak expression during hair follicle initiation (R^2^ = 0.89) D. Rat Xedar expression with fitted second-order polynomial demonstrating stable expression during scale development (R^2^ = 0.75)

**Figure 3:**
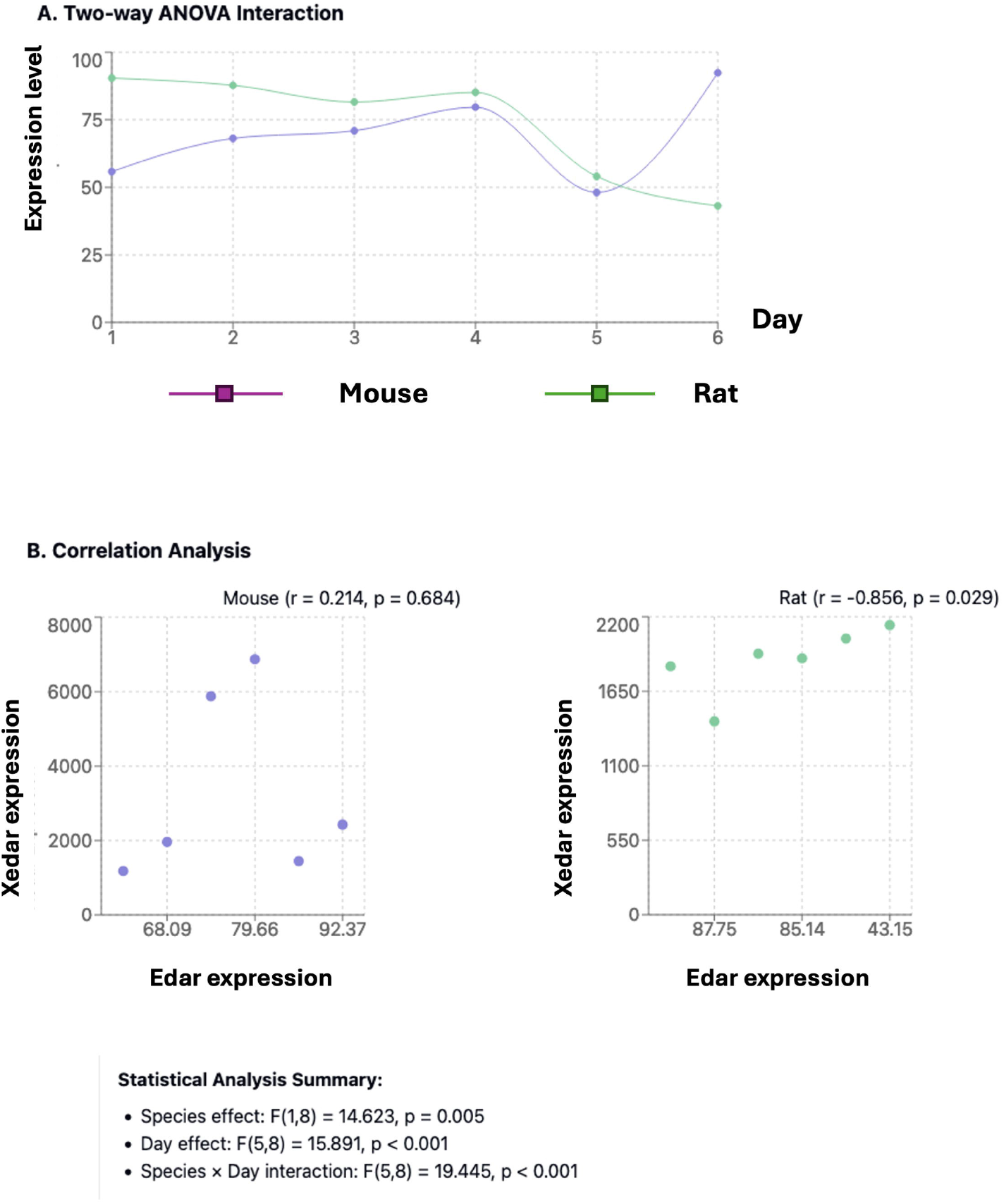
Statistical analysis of expression patterns A. Two-way ANOVA interaction plot showing species-specific temporal regulation of gene expression B. Correlation analysis between Edar and Xedar expression in mouse (r = 0.214, p = 0.684) and rat (r = −0.856, p = 0.029) C. Summary statistics showing significant differences in expression patterns between species and across developmental stages

**Figure 4:**
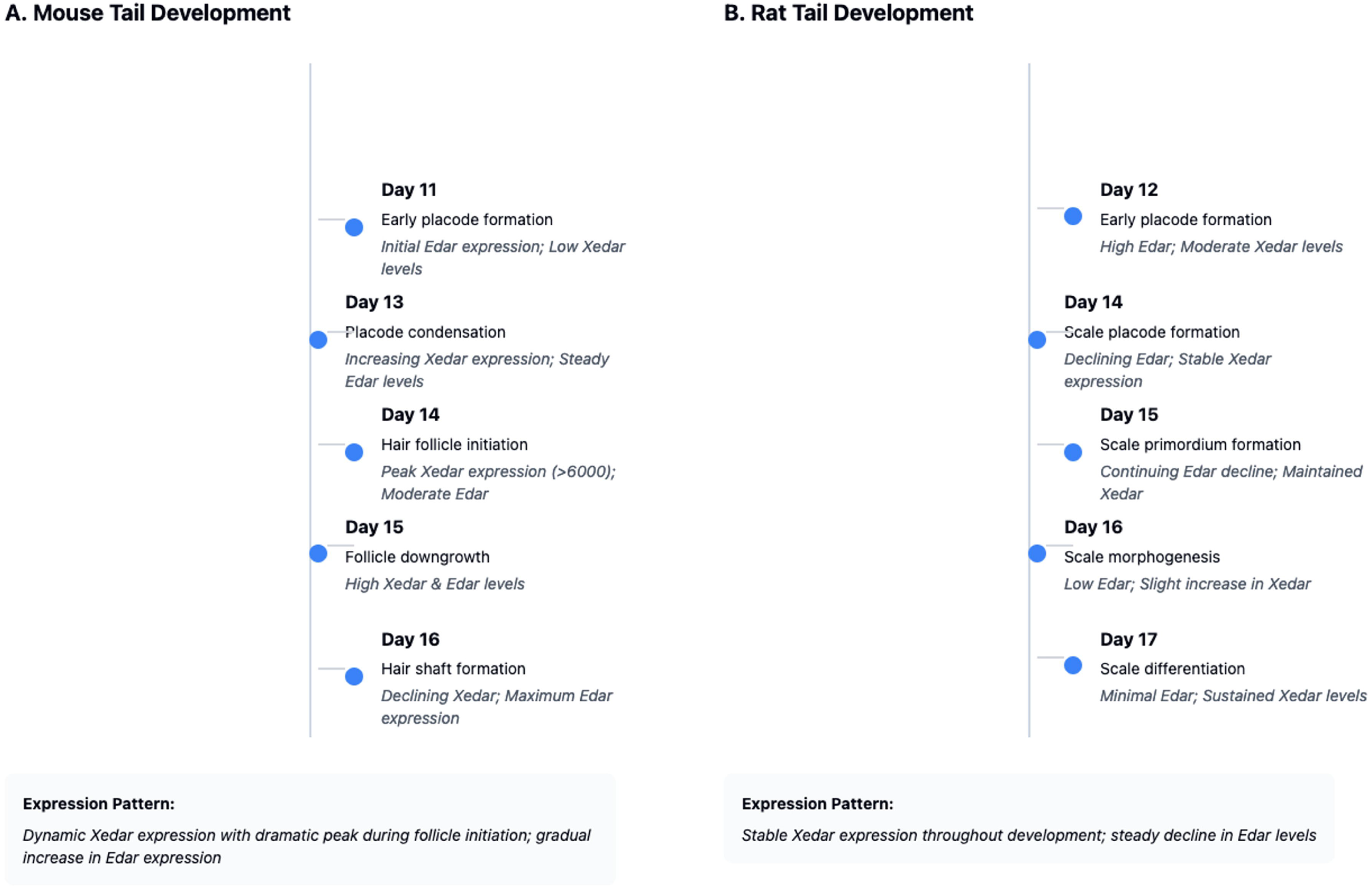
Developmental timeline of appendage formation and gene expression A. Mouse tail development timeline showing correlation between Edar/Xedar expression and critical stages of hair follicle formation (days 11-16) B. Rat tail development timeline showing relationship between gene expression and scale formation (days 12-17) Key morphological events and corresponding gene expression patterns are indicated for each developmental stage.

## Supporting information

Real-time PCR results data

## Author Contributions Statement

S.A.W. conceived and designed the study, performed the experiments, analyzed the data, wrote the manuscript, and approved the final version.

## Data Availability Statement

All data and protocols described in this manuscript are freely available on Figshare (DOI: will be provided upon acceptance) under a CC-BY 4.0 license.

## Accessory files

CSV file with Edar, Xedar, β-actin results – Figshare repository DOI: 10.6084/m9.figshare.27927177

## Scripts

R scripts used for analyses – Github repository https://github.com/SlawomirWisniewski73/edar-xedar-expression-analysis

## References

1. Bilgic, S. N., Domaniku, A., Toledo, B., Agca, S., Weber, B. Z. C., Arabaci, D. H., Ozornek, Z., Lause, P., Thissen, J.-P., Loumaye, A. and Kir, S. (2023), ‘EDA2R-NIK signalling promotes muscle atrophy linked to cancer cachexia.’, Nature 617, 827--834.

2. Bustin SA, et al. (2009) The MIQE guidelines: minimum information for publication of quantitative real-time PCR experiments. Clinical Chemistry 55(4): 611–622.

3. Chang, S. H., Jobling, S., Brennan, K. and Headon, D. J. (2009), ‘Enhanced Edar signalling has pleiotropic effects on craniofacial and cutaneous glands.’, PloS one 4, e7591.

4. Cooper G., Schiller A.L. (1975) Anatomy of the Guinea Pig. Harvard University Press, Cambridge.

5. Dhouailly D. (2009) A new scenario for the evolutionary origin of hair, feather, and avian scales. Journal of Anatomy, 214: 587–606.

6. Drew, C. F., Lin, C. M., Jiang, T. X., Blunt, G., Mou, C., Chuong, C. M. and Headon, D. J. (2007), ‘The Edar subfamily in feather placode formation.’, Developmental biology 305, 232--245.

7. Glover, J. D., Sudderick, Z. R., Shih, B. B.-J., Batho-Samblas, C., Charlton, L., Krause, A. L., Anderson, C., Riddell, J., Balic, A., Li, J., Klika, V., Woolley, T. E., Gaffney, E. A., Corsinotti, A., Anderson, R. A., Johnston, L. J., Brown, S. J., Wang, S., Chen, Y., Crichton, M. L. and Headon, D. J. (2023), ‘The developmental basis of fingerprint pattern formation and variation.’, Cell 186, 940--956.e20.

8. Held LI Jr. (2010) Quirks of Human Anatomy: An Evo-Devo Look at the Human Body. Cambridge University Press.

9. Kowalczyk-Quintas, C., Schuepbach-Mallepell, S., Willen, L., Smith, T. K., Huttner, K., Kirby, N., Headon, D. J. and Schneider, P. (2015), ‘Pharmacological stimulation of Edar signaling in the adult enhances sebaceous gland size and function.’, The Journal of investigative dermatology 135, 359--368.

10. Kozera B, Rapacz M (2013) Reference genes in real-time PCR. Journal of Applied Genetics 54(4): 391–406.

11. Lefebvre, S. and Mikkola, M. L. (2014), ‘Ectodysplasin research–where to next?’, Seminars in immunology 26, 220--228.

12. Lindfors, P. H., Voutilainen, M. and Mikkola, M. L. (2013), ‘Ectodysplasin/NF-κB signaling in embryonic mammary gland development.’, Journal of mammary gland biology and neoplasia 18, 165--169.

13. Livak KJ, Schmittgen TD (2001) Analysis of relative gene expression data using real-time quantitative PCR and the 2(-Delta Delta C(T)) Method. Methods 25(4): 402–408.

14. Mikkola, M. L. (2011), ‘The Edar subfamily in hair and exocrine gland development.’ Advances in experimental medicine and biology 691, 23--33.

15. Naito, A., Yoshida, H., Nishioka, E., Satoh, M., Azuma, S., Yamamoto, T., Nishikawa, S.-i. and Inoue, J.-i. (2002), ‘TRAF6-deficient mice display hypohidrotic ectodermal dysplasia.’, PNAS 99, 8766--8771.

16. Pantalacci, S., Chaumot, A., Benoît, G., Sadier, A., Delsuc, F., Douzery, E. J. P. and Laudet, V. (2008), ‘Conserved features and evolutionary shifts of the EDA signaling pathway involved in vertebrate skin appendage development.’, Molecular biology and evolution 25, 912--928.

17. Podzus, J., Kowalczyk-Quintas, C., Schuepbach-Mallepell, S., Willen, L., Staehlin, G., Vigolo, M., Tardivel, A., Headon, D., Kirby, N., Mikkola, M. L., Schneider, H. and Schneider, P. (2017), ‘Ectodysplasin A in Biological Fluids and Diagnosis of Ectodermal Dysplasia.’, Journal of dental research 96, 217--224.

18. Rieu I, Powers SJ (2009) Real-time quantitative RT-PCR: design, calculations, and statistics. The Plant Cell 21(4): 1031–1033.

19. Sadier, A., Twarogowska, M., Steklikova, K., Hayden, L., Lambert, A., Schneider, P., Laudet, V., Hovorakova, M., Calvez, V. and Pantalacci, S. (2019), ‘Modeling Edar expression reveals the hidden dynamics of tooth signaling center patterning.’, PLoS biology 17, e3000064.

20. Sadier, A., Viriot, L., Pantalacci, S. and Laudet, V. (2014), ‘The ectodysplasin pathway: from diseases to adaptations.’, Trends in genetics : TIG 30, 24--31.

21. Spearman, R.I.C. (1971) The integumentary morphology of the laboratory rat. Journal of Anatomy, 108: 13–29.

22. Taylor S, et al. (2010) A practical approach to RT-qPCR—Publishing data that conform to the MIQE guidelines. Methods 50(4): S1–S5.

23. Wark, A. R., Aldea, D., Tomizawa, R. R., Kokalari, B., Warder, B. and Kamberov, Y. G. (2023), ‘Ectodysplasin Signaling through XEDAR Is Required for Mammary Gland Morphogenesis.’, The Journal of investigative dermatology 143, 1529--1537.e2.

24. Wisniewski SA (2024) “Widespread expression patterns of Edar and Xedar during mouse and rat embryonic development-an in-situ hybridization study, DOI 10.6084/m9.figshare.27927180

25. Wu P, Hou L, Plikus M, Hughes M, Scehnet J, Suksaweang S, Widelitz R, Jiang TX, Chuong CM. (2004) Evo-Devo of amniote integuments and appendages. Int J Dev Biol. 48: 249–270.

26. Yan M, Wang LC, Hymowitz SG, Schilbach S, Lee J, Goddard A, de Vos AM, Gao WQ, Dixit VM. Two-amino acid molecular switch in an epithelial morphogen that regulates binding to two distinct receptors. Science. 2000 Oct 20;290(5491):523–7.

